# Computational conjugate adaptive optics for longitudinal through-skull imaging of cortical myelin

**DOI:** 10.1101/2022.05.18.492378

**Authors:** Yongwoo Kwon, Jin Hee Hong, Seokchan Yoon, Sungsam Kang, Hojun Lee, Yonghyeon Jo, Ki Hean Kim, Wonshik Choi

**Author notes:** These authors contributed equally to this work (Yongwoo Kwon, Jin Hee Hong, Seokchan Yoon).

## Abstract

We present a 1.3-μm reflection matrix microscope and computational conjugate adaptive optics algorithm for label-free longitudinal imaging of cortical myelin through an intact mouse skull. The myelination processes of the same mice were observed from 3 to 10 postnatal weeks down to cortical layer 4 with a near-diffraction-limited resolution of 0.79 µm. Our system will expedite the investigations on the role of myelination in learning, memory, and brain disorders.

## Main text

Myelin is a complex cellular structure, in which a lipid-rich multilayered membrane tightly wraps around the axons to provide electrical insulation and metabolic support. Myelin enables saltatory axonal conduction that substantially increases the neural processing speed and energy efficiency^1^. Myelin is mostly concentrated in the white matter in the central nervous system, as well as widespread in the gray matter. In particular, the myelin patterns are known to be dynamic in the mouse cortex with aging and learning^2,3^. It is also proposed that the myelination of pyramidal neurons located deep in the cortical layer can be an important substrate for neuroplasticity^4^. These results suggest that myelin could play an important role in higher brain functions, such as learning and memory. To further investigate myelin-related physiology and brain disorder, it is critical to observe the myelination processes from the early developmental stages and for a long period of time with minimal impairments to the subjects.

The longitudinal imaging of cortical myelin in live animals requires both high resolution and deep-tissue imaging capability because the cortical myelin is very thin, approximately 1 µm in diameter, and it is distributed from the shallow to deep layers of the brain cortex. Fluorescence imaging modalities relying on either external dye injection or genetic labeling have been used to visualize myelinating cells^5^. However, exogenous dyes cannot penetrate to a sufficient depth, and genetic labeling can visualize only a small fraction of mature myelination owing to its partial expression in a small population of oligodendrocytes. Label-free imaging modalities, such as optical coherence microscopy (OCM)^6^, spectral confocal reflectance microscopy^7^, third-harmonic generation microscopy^8^, and coherent Raman imaging^9^ have drawn special attention as they are free from these issues. However, all these methods require either the removal of the skull for the implantation of a cranial window or thinning of the skull to secure clear optical access. The skull opening and thinning procedures cause mechanical stress during surgery, leading to the activation of microglia and astrocytes^10^. They can particularly be disruptive in young subjects, thereby hampering the investigation of the early developmental stages. Recently, we reported a label-free reflectance imaging method, namely the laser-scanning reflection-matrix microscopy, with the source wavelength of 0.9 µm^11^, capable of imaging the myelin in cortex layer 1 of the mouse brain by correcting cranial aberrations. Due to the strong scattering and complex aberration by the mouse skull, the imaging fidelity was not high enough to reach cortical layer 4 and allow a longitudinal study.

Here, we report the longitudinal imaging of cortical myelin through an intact mouse skull to the depth of cortical layer 4 (∼ 650 µm below the dura matter). We constructed a high-speed reflection matrix microscope (RMM) using a light source with a wavelength of 1.3 µm, instead of 0.9 µm, for reducing tissue scattering and aberration^12,13^. Furthermore, we developed a computational conjugate adaptive optics (CC-AO) algorithm, for the first time, to optimally compensate for the skull aberrations. Compared with the hardware conjugate AOs^14-16^, this newly developed algorithm is flexible in the choice of the conjugate plane in the post-processing steps and can remove much more complex aberrations. These combined hardware and software improvements substantially raised the fidelity of the through-skull imaging, making it possible to visualize cortical layer 4 with near-diffraction-limited spatial resolution. We conducted label-free reflectance imaging of the individual myelin segments in the same area in the somatosensory cortex of the same mouse from its weaning stages (postnatal days 23–28 (P23–P28)) to its adulthood (P56–P70). In doing so, we quantified the growth rate of the myelination processes in cortical layer 1 and observed the emergence of new myelin segments in cortical layer 4 with aging.

The backbone of the 1.3-µm RMM system is an interferometric confocal microscope, but we replaced the confocal pinhole and photodetector with a camera to measure the phase and amplitude of all the elastic backscattering signals from the sample (see Supplementary Fig.1 for detailed experimental setup). A wavelength-tunable pulsed laser (INSIGHT X3, Spectra physics) with a bandwidth of 19 nm at the wavelength of 1.3 μm was used as the light source to provide a coherence gating window (or time-gating window) of 25 μm. The output beam from the laser was delivered to the sample via two galvanometer scanning mirrors for raster scanning with a focused illumination at the sample plane. A long-working-distance objective lens (XLPLN25XWMP2, Olympus, 25×, 1.05 NA) was used to ensure a high spatial resolving power (diffraction limit: 0.65 µm) for obtaining images from areas deep within the tissues. The backscattering signal from the sample was captured by the same objective lens and then delivered to the camera after being descanned by the same scanning mirrors. A fast InGaAs camera (Cheetah 800, Xenics, 6.8 kHz framerate) was placed at a plane conjugate to the sample to record the interference image formed by the elastic backscattering signal from the sample and the reference beams. The off-axis interferogram was processed to obtain the phase and amplitude maps of the sample waves for each illumination position. Unlike our previous setup (wavelength: 0.9 µm), the back-reflection noise from various optical components in the present setup was significantly large owing to the relatively low quality of the anti-reflection coating. A combination of a linear polarizer and a quarter-wave plate, centered at a wavelength of 1.31 µm, was installed to minimize the stray reflections from the optics. Our system can also be used to record confocal fluorescence and second harmonic generation (SHG) images by installing a dichroic mirror and a photomultiplier tube (H5784-20, Hamamatsu Photonics). The sample stage, scanning mirrors, data acquisition board, and camera were controlled via MATLAB.

In the data acquisition session, we recorded a set of complex-field maps while scanning the focus of illumination. The field of detection at the camera for each focused illumination was typically 25 × 25 μm^2^, and it took 4 s to scan the focus across a field of illumination of 100 × 100 μm^2^ with a scanning step size of the diffraction limit. A time-gated reflection matrix was constructed using the recorded complex-field maps, and the CC-AO algorithm was applied to the reflection matrix to reconstruct the aberration-free object image (see Methods and Supplementary section 2 for details). In our previous study, we employed an algorithm, called a closed-loop accumulation of single scattering (CLASS), to deal with the aberrations in the pupil plane^17^. This is well-suited for samples whose aberration sources are far from the sample plane. In the case of the through-skull imaging, the mouse skull is in proximity to the sample plane, causing a position-dependent aberration on the basis of the pupil correction. Therefore, the aberrations must be addressed for each small subregion (approximately 15 × 15 μm^2^ in the case of through-skull imaging), which undermine the fidelity of the aberration correction. Instead, we consider that the aberration layer is located at a plane conjugate to the skull and computationally change the basis of the reflection matrix from the sample plane to the plane where the skull is located. This made it possible to compensate for the skull-induced aberrations over the field of view as large as 80 × 80 μm^2^ with an enhanced image reconstruction fidelity and quality (see Supplementary Figs. 3 and 4). This strategy has been employed in the hardware-based AO microscopy^14-16^, wherein the wavefront shaping device is placed at a plane conjugate to the skull. Using the reflection matrix, we could realize the same concept in computational post-processing. In hardware conjugate AO, the conjugate plane set by the wavefront shaping needs to be carefully matched to the skull during the experiment. Furthermore, time-consuming feedback optimization should be performed prior to the image acquisition. On the contrary, our CC-AO can freely choose the optimal conjugate plane after the data acquisition, and ingenious and complex aberration correction operations can be administrated computationally to cope with much more complex aberrations.

We initially performed the through-skull deep-brain imaging up to cortical layer 4 for a five-week-old mouse (Fig. 1, Supplementary movie 1). The SHG images were acquired to determine the skull thickness (∼100 μm) (Fig. 1a). The depth of dura matter was set to 0 μm, and the SHG signal was displayed in the depth between -100 μm to 0 μm. Then, the time-gated reflection matrices were recorded over a field of view of 160 × 160 μm^2^ and in the depth range from 0 μm to 650 μm beneath the dura at a depth interval of 3 μm. This depth range covers the cortical layers 1 to 4. In each recording, the reference beam path was adjusted based on the average refractive index of the mouse brain tissue. The full recorded field of view was divided into 5 × 5 subregions, and the CC-AO algorithm was applied to each submatrix to identify the aberrations at the corresponding subregion (Fig. 1d). To minimize the image contrast mismatch at the edges of the subregions, the area of individual subregions was chosen to be 64 × 64 μm^2^, and then the reconstructed images were overlapped with each other in the ratio of 50 %. The subregion area of 64 × 64 μm^2^ processed by the CC-AO algorithm was much larger than the isoplanatic patch size (15 × 15 μm^2^) used in the previous pupil-plane CLASS (Supplementary Fig. 3). It should be noted that the aberration maps indicate the phase delay induced by the skull at the conjugate plane. Considering the illumination/detection geometry, the area of the aberration map becomes larger as the imaging depth increases (see the scale bars in Fig. 1d). The aberration-corrected images from all the subregions were merged to form a full-field image at each depth, and a 3D-rendered volumetric image was reconstructed from all the depth-dependent images (Fig. 1a). For a few representative depths, the aberration-corrected images were displayed using a maximum intensity projection (MIP) over a depth range of ±15 μm (Fig. 1c, Supplementary movies 2 and 3). As a point of reference, the conventional OCM images obtained from the same reflection matrices are shown in Fig. 1b.

**Figure 1.**
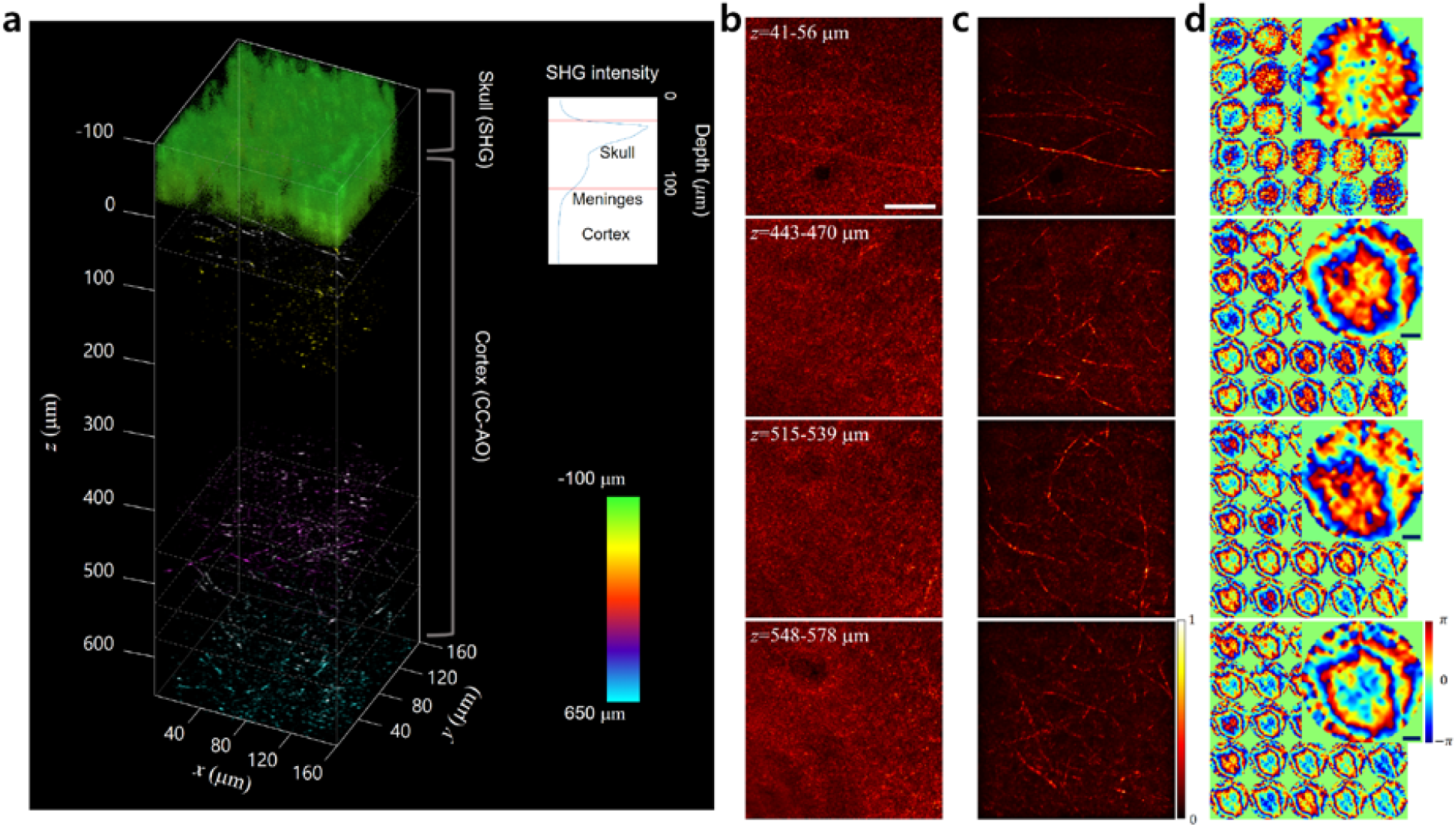
In vivo reflectance images of a mouse brain down to cortical layer 4 through an intact skull. **a**, 3D rendered through-skull image. The SHG images were used to visualize the mouse skull (colored in green). The depth-dependent intensity was plotted to estimate the skull thickness (inset). The depth of the dura matter was set as *z* = 0 µm. Cortical layer 1 (up to ∼100 μm) and cortical layer 4 (400–650 μm) were mainly imaged since the myelinated axons are distributed laterally at these layers. Color map, depth. **b**, Conventional OCM images at representative depths displayed with gray-dotted boxes in **a**. Scale bar, 40 μm. **c**, Aberration-corrected images at the same depths as those shown in **b**. Individual images are MIPs in the depth ranges of 41–56, 443–470, 515– 539, and 548–578 μm. Each image was normalized by its maximum intensity. Color bar, normalized intensity. **d**, Aberration maps of the subregions in **c**. Color bar, phase in radians. Zoomed-in images show representative aberration maps at the conjugate planes. Scale bars, 100 μm.

Due to severe skull-induced aberrations, the conventional OCM displayed the microstructures at cortical layer 1 (0–100 µm) only vaguely and completely lost its resolving power as the depth was further increased. After compensating for the aberrations, numerous myelin fibers up to cortical layer 4 were clearly resolved (Fig. 1c). We proved that the fibril structures in the reflectance image were myelinated axons, via a separate *ex vivo* fluoromyelin-stained fluorescence imaging (Supplementary Fig. 5). The thinnest myelin observed at a depth of 518 µm beneath the dura (cortical layer 4) was 0.79 μm in diameter, which is close to the diffraction-limited resolution (Supplementary Fig. 6). We observed that the signal enhancement after the aberration correction grew exponentially with depth (Supplementary Fig. 7c). The signal enhancement at cortical layer 4 was approximately 250 times. This trend is consistent with the Strehl ratio obtained through the point spread function (PSF) analysis of the aberration maps in Fig. 1d (Supplementary Fig. 7b). The PSF width was broadened by 44 times relative to the diffraction limit at cortical layer 4, supporting the difficulty in reaching cortical layer 4 via the intact skull.

We conducted a longitudinal observation of a mouse brain up to cortical layer 4 through the intact skull, from its weaning period to the adult stage (Fig. 2). We could start the investigation at an early stage, as young as three weeks after the birth, because the intact skull imaging is a non-invasive technique that does negligible harm to the subject. The same region of the mouse brain was repeatedly imaged by keeping track of the blood vessel structures beneath the skull. In each imaging session, a volumetric image was acquired from the surface of the skull up to a depth of 650 µm below the dura. However, the analysis was mainly focused on cortical layers 1 and 4, where the axons stretch laterally. Figure 2a shows the MIP images of cortical layer 1 recorded at P26, P33, and P47 over a field of view of 400 × 400 μm^2^ and in a depth range between ∼40 µm to ∼90 µm below the dura. Notably, different depth ranges were chosen for comparing the myelin at different ages because of the growth of the mouse skull and brain. A drastic increase in the density of myelin was observed, especially at a very early stage of the lifetime of the mice. The high-resolution images allowed us to trace the growth of the individual myelination processes. For example, the yellow and white arrowheads in Fig. 2a indicate the newly emerged myelin segments that did not appear in the preceding stages.

**Figure 2.**
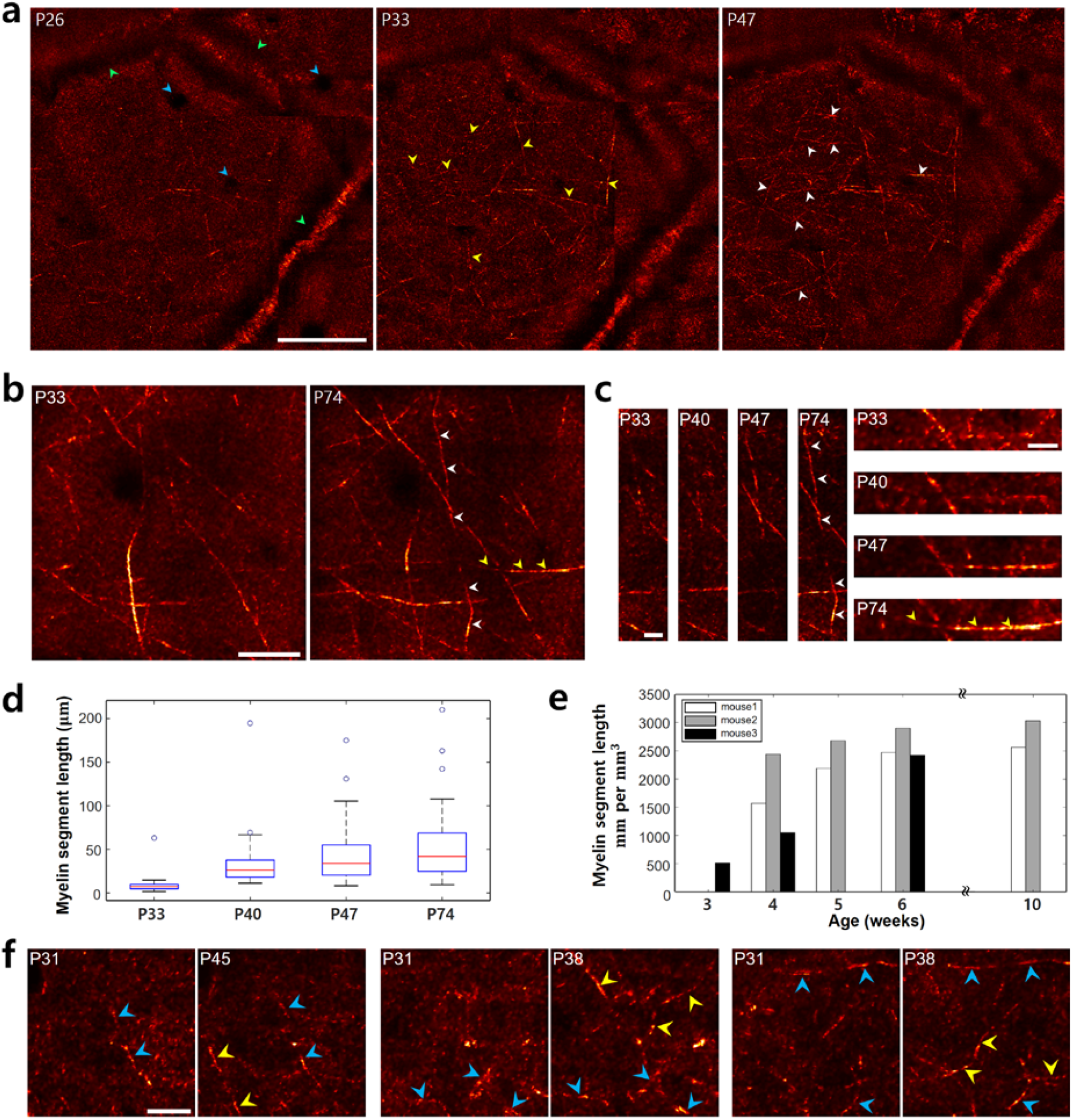
Longitudinal observation of the myelination processes up to cortical layer 4 through the intact skull. **a**, Aberration-corrected images at cortical layer 1 of the same mouse recorded at P26, P33, and P47. Each image is an MIP across a 40-μm depth range. The yellow and white arrowheads mark the newly emerged myelin segments. Blood vessels oriented along the lateral and axial directions are marked with green and blue arrowheads, respectively. Scale bar, 100 μm. **b**, Aberration-corrected images at cortical layer 1 of a different mouse from **a**, taken at P33 and P74. Each image is an MIP across a 20-μm depth range. Scale bar, 30 μm. **c**, Zoomed-in images of the myelin segments marked with white and yellow arrowheads shown in **b**. Images are shown at P33 and P74 as well as at P40 and P47. Scale bars, 10 μm. **d**, Myelin segment lengths measured from the images in **b**. Red lines indicate median values, and open dots show the outliers. **e**, Total myelin segment length per unit volume derived from three different mice with age. **f**, Aberration-corrected images on two different postnatal days at cortical layer 4. Three examples are shown. These individual pairs were MIPs over the depths of 405–435, 455– 485, and 510–540 μm (from the left). The newly emerged myelin segments were marked with yellow arrowheads, whereas the remaining unchanged ones with age were marked with blue arrowheads. Scale bar, 20 μm.

A similar progression of the myelination processes was consistently observed in another mouse (Figs. 2b and c). In this case, the growth of myelination was monitored at P33, P40, P47, and P74. In Fig. 2b, the myelin segments indicated by the white and yellow arrowheads at P74 are absent at P33. Figure 2c shows the gradual elongation of these myelin segments. We measured the lengths of individual myelin segments derived from the MIP images shown in Fig. 2b (Fig. 2d). The median values of myelin segment length increased with age. In addition, we measured the sum of all the myelin segment lengths per unit volume in cortical layer 1 for three different mice (Fig. 2e). Although there were slight variations among different mice and observation areas, substantial growth was observed, especially during the postnatal weeks 3–5. The system’s capability of high-resolution deep brain imaging enabled us to investigate the myelination processes even at layer 4 (Fig. 2f). Like cortical layer 1, we observed the emergence of new myelin segments (yellow arrowheads) as well as the persistence of the existing segments (blue arrowheads). In fact, this is the first label-free observation of individual myelin segments at layer 4 through an intact skull. Notably, the image contrast of the myelin segments tended to increase over time (for example, see Fig. 2c). Since the lipid constituting the myelin is the main source of the intrinsic reflectance contrast, this observation implies the gradual thickening of the myelin. In addition to the myelin, blood vessels stretching laterally (green arrowheads in Fig. 2a) and axially (blue arrowheads in Fig. 2a) were also visible.

In summary, our 1.3-µm RMM system and image reconstruction algorithm enabled the tracing of individual myelin segments in the cortical brain, without cranial surgery. This allowed us to investigate the myelination processes at the early developmental stages that are prone to surgical side effects. In addition to the suppression of scattering and aberration by a longer wavelength source, we found that the newly developed CC-AO algorithm played a critical role. In the previous pupil AO, the full-field image was divided into small subregions with sizes comparable to the isoplanatic patch (15 × 15 μm^2^) within which the aberration is identical. The fidelity of aberration correction is determined by the competition between the coherent addition of signals and the incoherent addition of multiple scattering noise, which favors a large isoplanatic patch. Therefore, a small isoplanatic patch size results in the reduced fidelity of the aberration correction, especially when the skull thickness increases with aging. There is also degeneracy in the tilt and defocus in the pupil AO, which can give rise to the lateral and axial shifts of reconstructed images in the individual patches. This makes it difficult to trace myelin segments in the volume and causes the blur of the MIP images (see Supplementary Fig. 4). In contrast, the CC-AO algorithm could provide reliable aberration correction over the wide field of view, whose size is mainly determined by the computer memory. In our present study, this field of view was approximately 80 × 80 μm^2^, i.e., ∼30 times the isoplanatic patch (see Supplementary Fig. 3). It should be noted that the implementation of the CC-AO was possible because of the recording of the reflection matrix, which is a replica of the real optical system. Using the reflection matrix, one can freely choose the basis of illumination/detection from the pupil plane to any plane suitable for correcting the skull aberrations in post-processing. Our system can serve as a potential platform for investigating the early development of learning and memory associated with myelination^18^. It can also be used for understanding myelin-related disorders, such as multiple sclerosis and leukodystrophies^19^, and developing their treatment strategies. Technically, our system can be used as a wavefront sensing AO for nonlinear deep-brain imaging modalities, such as two- or three-photon microscopy, to recover their resolving power in the through-skull imaging^20^. This will facilitate longitudinal deep brain imaging of many interesting neuronal structures, such as axons and dendrites, as well as non-neuronal structures, such as oligodendrocytes and microglia.

## Methods

### Computational conjugate adaptive optics algorithm

The reflection matrix ***R*** is measured on the basis of a sample plane set by the objective focus, whose lateral coordinates are **r** = (*x, y*). The matrix element describes an electric field at a detector point conjugate to **r**_o_ = (*x*_o_, *y*_o_) under the illumination of a point source at **r**_in_ = (*x*_in_, *y*_in_). For the conjugate AO, we change the input and output bases to a plane **w** = (*u, v*) located at a distance *z* from the object plane, where the skull is located. The matrix element of the conjugate reflection matrix ***R***_con_ is the electric field *E*(**w**_o_; **w**_in_) at a detector point conjugate to **w**_o_ = (*u*_o_, *v*_o_) under the illumination at a point **w**_in_ = (*u*_in_, *v*_in_). Under Fresnel approximation, it is written as

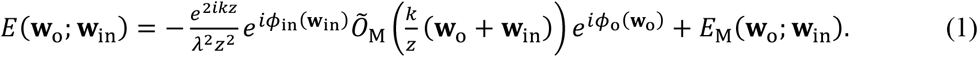

Here, 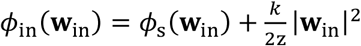 and 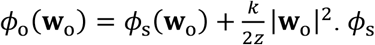 is the aberration map of the skull at the conjugate plane (*k* = 2*π*/*λ* with *λ* the wavelength of the light source). *Õ*_M_ is the 2D Fourier transform of 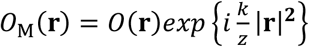, where *0*(**r**) is the amplitude reflectance of the target object at the sample plane (see Supplementary section 2 for the detailed derivation of Eq. (1)). *E*_M_(**w**_o_; **w**_in_) is the multiple scattering noise from the skull and brain tissues. Equation (1) resembles 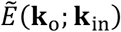, the Fourier transform of *E*(**r**_o_; **r**_in_) with respect to **r**_o_ and **r**_in_, on the basis of pupil aberrations^17^.

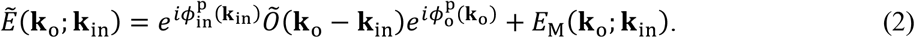

Here 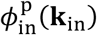 and 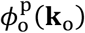 are the pupil aberrations. We decomposed *ϕ*_in_(**w**_in_), *ϕ*_o_(**w**_o_), and *Õ*_M_ from ***R***_con_ by modifying the algorithm that decomposes 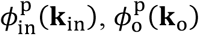 and *Õ* from ***R***. The skull aberration *ϕ*_s_, and the reflectance image *0*(**r**) are obtained from *ϕ*_in_(**w**_in_) (or *ϕ*_o_(**w**_o_)) and *Õ*_M_ after removing the quadratic phase terms.

### Animal preparation for the longitudinal through-skull imaging

We employed the procedure outlined in previous reports^11,20^, with slight modifications. Three- to four-week-old C57BL/6 mice (P21-P28; body weight: 9-14 g) were anesthetized with isoflurane (2 % in oxygen for induction, and 1.0-1.5 % in oxygen for surgery to maintain a breathing frequency of around 1 Hz). The body temperature was kept at 37-38 °C by a heat blanket, and the eyes were protected with an eye ointment during the surgery and imaging. Dexamethasone (2 mg/kg) was administrated via subcutaneous injection to minimize swelling at the surgery site. The hair and scalp were removed to expose the bregma and lambda, and both parietal plates of the skull. The connective tissue remaining on the skull was gently removed with sterile saline. After the saline that covered the skull was completely removed, a sterile coverslip of 5-mm diameter was attached to the center of the parietal bone using an ultraviolet-curable glue. A custom-made metal plate was attached to the skull with cyanoacrylate for head fixation during the *in vivo* imaging, and the exposed part of the skull was covered with dental cement. Finally, the window was covered by a biocompatible silicone sealant (Kwik-Cast, World Precision Instruments) for protection until the imaging was performed. For imaging, the mice were anesthetized with isoflurane (1.2-1.5 % in oxygen to maintain a breathing frequency of around 1.5 Hz) and placed on a 3D motorized stage heated by a heat blanket at 37-38 °C. The mice were weighed each time they were imaged during the experiment. All the animal-related procedures were approved by the Korea University Institutional Animal Care and Use Committee (KUIACUC-2019-0024).

### Measuring myelin segments

The aberration-corrected images were acquired at a depth interval of 3 µm from the cortical surface to a depth of ∼100 µm in cortical layer 1 of all the mice. The aberration-corrected images acquired using MATLAB were converted to TIFF images. For measuring the myelin segment lengths, these images were transferred to ImageJ. The image brightness and contrast levels were adjusted for clarity, and the size of the image was defined with “Set Scale.” Myelin sheaths within the field of view were drawn by a few investigators, and the lengths of the myelin segments were measured automatically.

## Acknowledgments

This work is supported by the Institute for Basic Science (IBS-R023-D1) and National Research Foundation (NRF) of Korea grant (NRF-2020R1A2C3009309).

## Author contributions

Y.K. and S.Y. designed and constructed an experimental setup along with H.L. J.H.H. designed mouse brain imaging sessions along with K.H.K. and undertook animal preparations. S.Y. and S.K. developed the image reconstruction algorithm. Y.K. and J.H.H. conducted experiments, data analysis, and image processing along with Y.J. Y.K., J.H.H., S.Y., and W.C. prepared the manuscript, and all authors contributed to finalizing the manuscript. W.C. supervised the project.

